# Population genomics of *Cryptococcus neoformans* var. *grubii* reveals new biogeographic relationships and finely maps hybridization

**DOI:** 10.1101/132894

**Authors:** Johanna Rhodes, Christopher A. Desjardins, Sean M. Sykes, Mathew A. Beale, Mathieu Vanhove, Sharadha Sakthikumar, Yuan Chen, Sharvari Gujja, Sakina Saif, Anuradha Chowdhary, Daniel John Lawson, Vinicius Ponzio, Arnaldo Lopes Colombo, Wieland Meyer, David M. Engelthaler, Ferry Hagen, Maria Teresa Illnait-Zaragozi, Alexandre Alanio, Jo-Marie Vreulink, Joseph Heitman, John R. Perfect, Anastasia Litvintseva, Tihana Bicanic, Thomas S. Harrison, Matthew C. Fisher, Christina A. Cuomo

## Abstract

*Cryptococcus neoformans* var. *grubii* is the causative agent of cryptococcal meningitis, a significant source of mortality in immunocompromised individuals, typically HIV/AIDS patients from developing countries. Despite the worldwide emergence of this ubiquitous infection, little is known about the global molecular epidemiology of this fungal pathogen. Here we sequence the genomes of 188 diverse isolates and characterized the major subdivisions, their relative diversity and the level of genetic exchange between them. While most isolates of *C. neoformans* var. *grubii* belong to one of three major lineages (VNI, VNII, and VNB), some haploid isolates show hybrid ancestry including some that appear to have recently interbred, based on the detection of large blocks of each ancestry across each chromosome. Many isolates display evidence of aneuploidy, which was detected for all chromosomes. In diploid isolates of *C. neoformans* var. *grubii (*serotype A/A) and of hybrids with *C. neoformans* var. *neoformans* (serotype A/D) such aneuploidies have resulted in loss of heterozygosity, where a chromosomal region is represented by the genotype of only one parental isolate. Phylogenetic and population genomic analyses of isolates from Brazil revealed that the previously ‘African’ VNB lineage occurs naturally in the South American environment. This suggests migration of the VNB lineage between Africa and South America prior to its diversification, supported by finding ancestral recombination events between isolates from different lineages and regions. The results provide evidence of substantial population structure, with all lineages showing multi-continental distributions demonstrating the highly dispersive nature of this pathogen.

**Author Summary:** *Cryptococcus neoformans* var. *grubii* is a human fungal pathogen of immunocompromised individuals that has global clinical impact, causing half a million deaths per year. Substantial genetic substructure exists for this pathogen, with two lineages found globally (VNI, VNII) whereas a third has appeared confined to sub-Saharan Africa (VNB). Here, we utilized genome sequencing of a large set of global isolates to examine the genetic diversity, hybridization, and biogeography of these lineages. We found that while the three major lineages are well separated, recombination between the lineages has occurred, notably resulting in hybrid isolates with segmented ancestry across the genome. In addition, we showed that isolates from South America are placed within the VNB lineage, formerly thought to be confined to Africa, and that there is phylogenetic separation between these geographies that substantially expands the diversity of these lineages. Our findings provide a new framework for further studies of the dynamics of natural populations of *C. neoformans* var. *grubii*.

## Introduction

The environmental basidiomycetous yeast *Cryptococcus neoformans* is capable of causing invasive fungal infections primarily in immunocompromised individuals. Meningitis is the most serious manifestation of cryptococcosis. The HIV/AIDS pandemic increased the population of these susceptible individuals and led to an increase in *C. neoformans* infection rates (Day, 2004). *C. neoformans* is the leading cause of mortality in HIV/AIDS patients worldwide, particularly in sub-Saharan Africa, where approximately half a million deaths occur annually (Park et al., 2009). While cryptococcal infection rates in HIV positive individuals have declined due to highly active antiretroviral therapy (HAART), new estimates continue to suggest there are more than 100,000 deaths/year (Rajasingham et al., 2017); recent data also suggests that the incidence of cryptococcosis has plateaued at a high number despite HAART availability. Furthermore, the increasing number of people living with other immunodeficiencies, including transplant and cancer patients, represents a growing population at risk for cryptococcosis (Maziarz and Perfect, 2016).

There are three major serotypes of *C. neoformans* distinguished by different capsular antigens, which include two separate varieties (*Cryptococcus neoformans* var. *grubii* and *Cryptococcus neoformans* var. *neoformans*, serotypes A and D respectively) and a hybrid between the two (serotype AD). While *C. neoformans* isolates are primarily haploid, diploid AD hybrid isolates consisting of both serotype A (*Cryptococcus neoformans* var. *grubii*) and serotype D (*Cryptococcus neoformans* var. *neoformans*) have been isolated from both clinical and environmental sources mostly in Europe (Cogliati, 2013; Desnos-Ollivier et al., 2015; Franzot et al., 1999). Serotype A isolates are the most common cause of infection, accounting for 95% of all *C. neoformans* infections globally (Casadevall and Perfect, 1998; Heitman et al., 2011). Genomes of serotype A and D isolates differ by 10-15% at the nucleotide level (Janbon et al., 2014; Kavanaugh et al., 2006; Loftus et al., 2005), and laboratory crosses of A and D isolates are possible but show reduced viability of meiotic spores (Lengeler et al., 2001; Vogan and Xu, 2014).

*Cryptococcus neoformans* var. *grubii* can be divided into three molecular types, or lineages: VNI, VNII and VNB (Litvintseva et al., 2006; Meyer et al., 1999, 2009). The VNI and VNII lineages are isolated globally, while the VNB lineage is predominantly located in sub-Saharan Africa (Litvintseva et al., 2006), although there is some evidence for VNB occurring in South America (Bovers et al., 2008; Ngamskulrungroj et al., 2009) and in the USA, Italy, and China in AD hybrid isolates (Litvintseva et al., 2007). Apart from clinical isolation, the VNI lineage is primarily associated with avian excreta (Lugarini et al., 2008; Nielsen et al., 2007) while the VNB lineage is found mostly in association with specific tree species predominantly mopane trees (Litvintseva and Mitchell, 2012; Litvintseva et al., 2011). These and recent studies have shown that VNI infections are associated with urbanized populations where an avian-associated reservoir, pigeon guano, is also found, while the VNB lineage is widely recovered in the African arboreal environment (Litvintseva et al., 2011; Vanhove et al., 2017).

Mating in *C. neoformans* occurs between cells of opposite mating types (*MAT***a** and *MAT*α) (Kwon-Chung, 1975, 1976), although unisexual mating can also occur (Lin et al., 2005). *MAT*α isolates are capable of unisexual mating both within and between the two serotypes (Lin et al., 2005, 2007), and recombination was shown to occur at similar levels in bisexual and unisexual mating in serotype D isolates (Desnos-Ollivier et al., 2015; Sun et al., 2014). Due to the rarity of *MAT***a** isolates of both serotypes in the environment (Lengeler et al., 2000a; Litvintseva et al., 2003; Viviani et al., 2001), unisexual mating may have evolved to enable meiotic recombination and genetic exchange between isolates. Several studies have found evidence of recombination within VNI, VNII, and VNB populations although not between these lineages (Bui et al., 2008; Litvintseva et al., 2003, 2005).

An additional level of genome diversity detected in *Cryptococcus neoformans* var. *grubii* includes the presence of cryptic diploid isolates and variation in the copy number of individual chromosomes or regions. Close to 8% of *Cryptococcus neoformans* var. *grubii* global isolates appear diploid; these isolates contain the *MAT*α locus and many appear autodiploid, thought to result either from endoreduplication or self-mating (Lin et al., 2009). While the vast majority of serotype A or D isolates appear haploid, individual chromosomes can be present at diploid or triploid levels (Hu et al., 2011). For chromosome 1, a specific advantage of aneuploidy is copy number amplification of the azole drug targets or efflux transporters, associated with drug resistance (Sionov et al., 2010). While the specific selective advantage of other chromosomal aneuploidies is unknown, same-sex mating of *MAT*α isolates generates aneuploid progeny at high frequency, some of which also exhibit azole resistance (Ni et al., 2013). Titan cells, polyploid yeast cells produced in the lung of infected animals, also generate aneuploid progeny under stress conditions (Gerstein et al., 2015).

Previous studies examining the global population structure of *Cryptococcus neoformans* var. *grubii* have used typing methods for a few genetic loci or focused on particular geographical regions or countries (Hiremath et al., 2008; Khayhan et al., 2013; Litvintseva et al., 2006; Oliveira et al., 2004). Recent approaches have applied whole genome sequencing (WGS) to trace the microevolution of *Cryptococcus*, identifying variation that occurs during the course of infection (Chen et al., 2017; Ormerod et al., 2013; Rhodes et al., 2017) or in the environment (Vanhove et al., 2017). Here, we use WGS of 188 isolates to provide a comprehensive view of the population variation between the three major lineages; the sequenced isolates were selected to represent the diversity of *Cryptococcus neoformans* var. *grubii* including each of the three major lineages and global geographical sampling. We identify contributions to genomic diversity generated through inter-lineage meiotic exchange to create haploid hybrids, generation of AD diploid hybrids, and regional copy number amplification. Furthermore, we finely analyze the phylogenetic relationships and trace the evolution of *Cng* at the global population level.

## Results

### Population subdivisions and detection of genetic hybrids

To examine the evolution of *C. neoformans* var. *grubii*, we sampled the population by sequencing the genomes of 188 isolates (Table 1, **Table S1**) representing each of the three major genetic subpopulations (VNI, VNII, and VNB) previously defined using multi-locus sequence typing (MLST) (Litvintseva et al., 2006; Meyer et al., 2009). These isolates are geographically diverse, originating from North America, South America, the Caribbean, Asia, Europe, and Africa (**Table S1**). The VNI global lineage is the most geographically diverse, whereas VNII is represented by a smaller number of locations and VNB appears most highly prevalent in southern Africa. For VNI and VNB, both clinical and environmental isolates were included, with 25 VNI isolates originating from avian guano or trees and 7 VNB isolates from trees or other environmental sources (**Table S1**). For each isolate we identified SNPs using GATK by aligning Illumina reads to the H99 reference genome assembly (Methods, (Janbon et al., 2014)). Whereas 164 isolates appeared haploid, 24 isolates were determined to be heterozygous diploids (Methods, Table 1) and analyzed separately. An initial phylogeny of the 164 haploid isolates separated the three lineages but intermediate placement of five isolates suggested the presence of hybrid haploid genotypes (**Figure S1**). As the phylogenetic placement of such hybrid isolates is complicated by recombination, we removed these isolates from the phylogenetic analysis and analyzed them using alternative approaches (see below).

**Table 1.** Properties of Sequenced isolates.

| Population | Isolates (#) | *MAT*α | *MATa* |
| --- | --- | --- | --- |
| VNI | 111 | 109 | 2 |
| VNII | 23 | 23 | 0 |
| VNB | 25 | 21 | 4 |
| VNI/VNB | 5 | 1 | 4 |
| VNII/VNB | 2 | 2 | 0 |

Diploid isolates
| Population | Isolates (#) | *MAT*α/*MA T*α | *MATa/MATa* | *MATa/MAT*α |
| --- | --- | --- | --- | --- |
| VNI/VNB | 1 | 1 | 0 | 0 |
| VNII/VNB | 2 | 2 | 0 | 0 |
| VNB/Cnn | 8 | 0 | 1 | 7 |
| VNB/Cng | 1 | 1 | 0 | 0 |

A phylogeny inferred from the SNPs for all non-hybrid isolates strongly supports the three major lineages of *C. neoformans* var. *grubii*: VNI, VNII, and VNB (Figure 1). Of these 159 isolates, only 6 (4%) contain the rare *MAT****a*** allele, including four VNB isolates (Bt63, Bt85, Bt206, and CCTP15) and two VNI isolates (125.91 and Bt130). Based on these whole genome SNP comparisons, none of these *MAT****a*** isolates appeared highly related to each other or to any *MAT*α isolate. The two VNI *MAT***a** isolates are well separated within this group, with Bt130 found in a subgroup of African isolates and 125.91 most closely related to a pair of isolates from Africa and North America (Figure 1). Phylogenetic analysis showed that VNB has the highest diversity between isolates, showing the longest tip branches compared to VNI or VNII. In addition, VNB consisted of two diverged subgroups, VNBI and VNBII, as suggested previously by MLST (Chen et al., 2015; Litvintseva et al., 2006, 2011) and genomic analysis (Desjardins et al., 2017; Vanhove et al., 2017).

**Figure 1.**
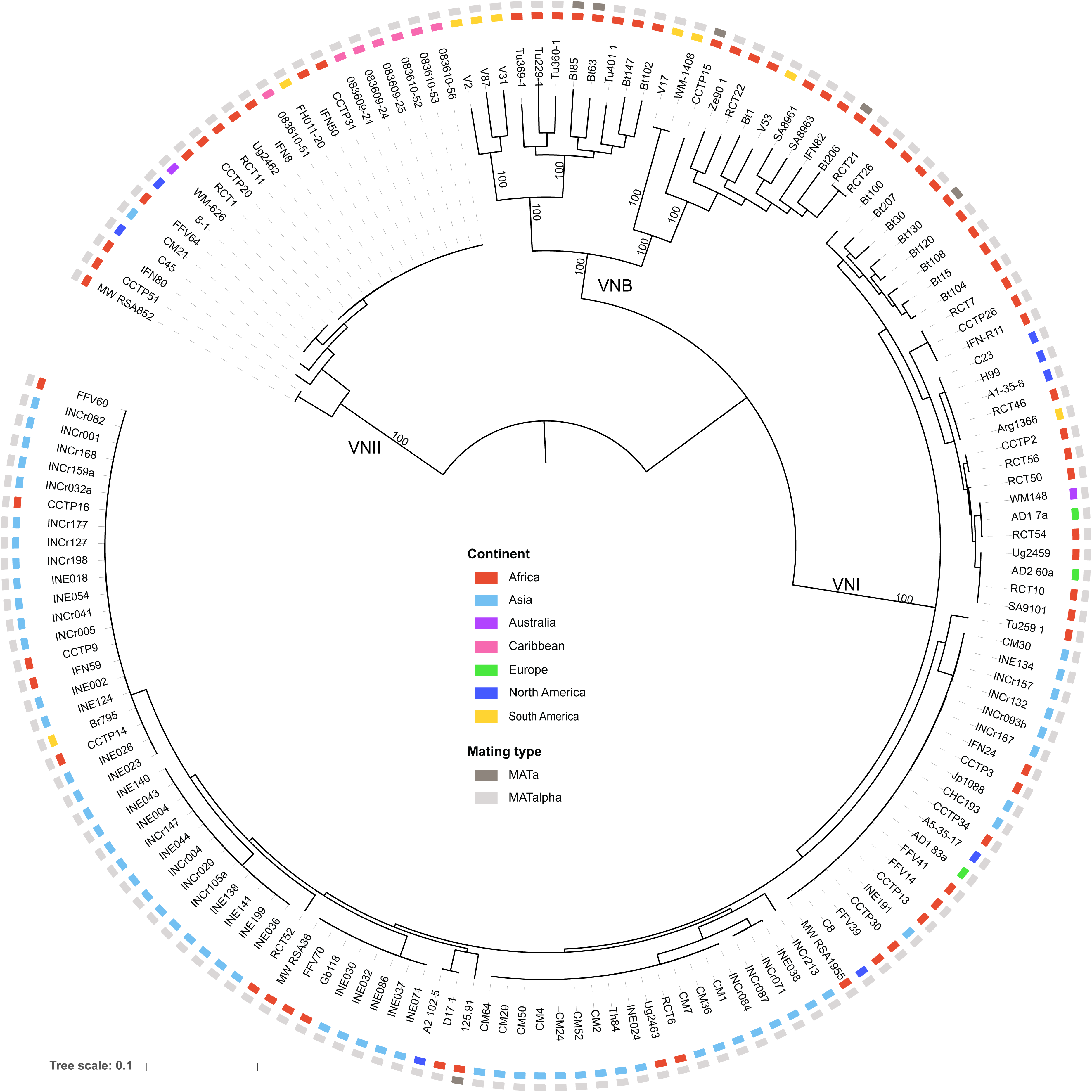
Phylogenetic analysis supports three major lineages of *C. neoformans* var. *grubii*. Using a set of 876,121 SNPs across the 159 non-hybrid isolates, the phylogenetic relationship was inferred using RAxML. The percentage of 1,000 bootstrap isolates that support each node is shown for major nodes with at least 90% support. For each isolate, the geographical site of isolation is noted by colored boxes.

To better understand the population structure of the three lineages and identify potential inter-lineage recombination, we compared results of two independent approaches. First, we used principle components analysis (PCA) to identify the major groups in the population using the SNP data. By comparing the SNP variants across isolates using PCA, we found there are three major clusters corresponding to the VNI, VNII, and VNB lineages (Figure 2). The five isolates that showed intermediate positions in phylogenetic analysis (**Figure S1**) also appeared at intermediate positions by PCA, placed between VNI and VNB. In addition, two isolates were separated from the VNII cluster and shifted towards the VNB cluster. All of these seven isolates were collected from southern Africa, and all had a clinical origin except isolate Ftc260-1, which was isolated from the environment (**Table S1**). Of the seven, two sets of isolates share nearly identical ancestry ratios and are closely linked on the phylogenetic tree. Isolates Bt131, Bt162, and Bt163 differed by an average of only 39 SNP positions; similarly CCTP51 and MW_RSA852 differed by 200 SNP positions, suggesting these five isolates are descended from two hybridization events. Therefore, four unique hybridization events were detected in total, three for VNI-VNB and one for VNII-VNB. While the basal branching VNB isolates from Brazil could suggest a hybrid ancestry, all appear to be uniformly VNB (>99% of sites).

**Figure 2.**
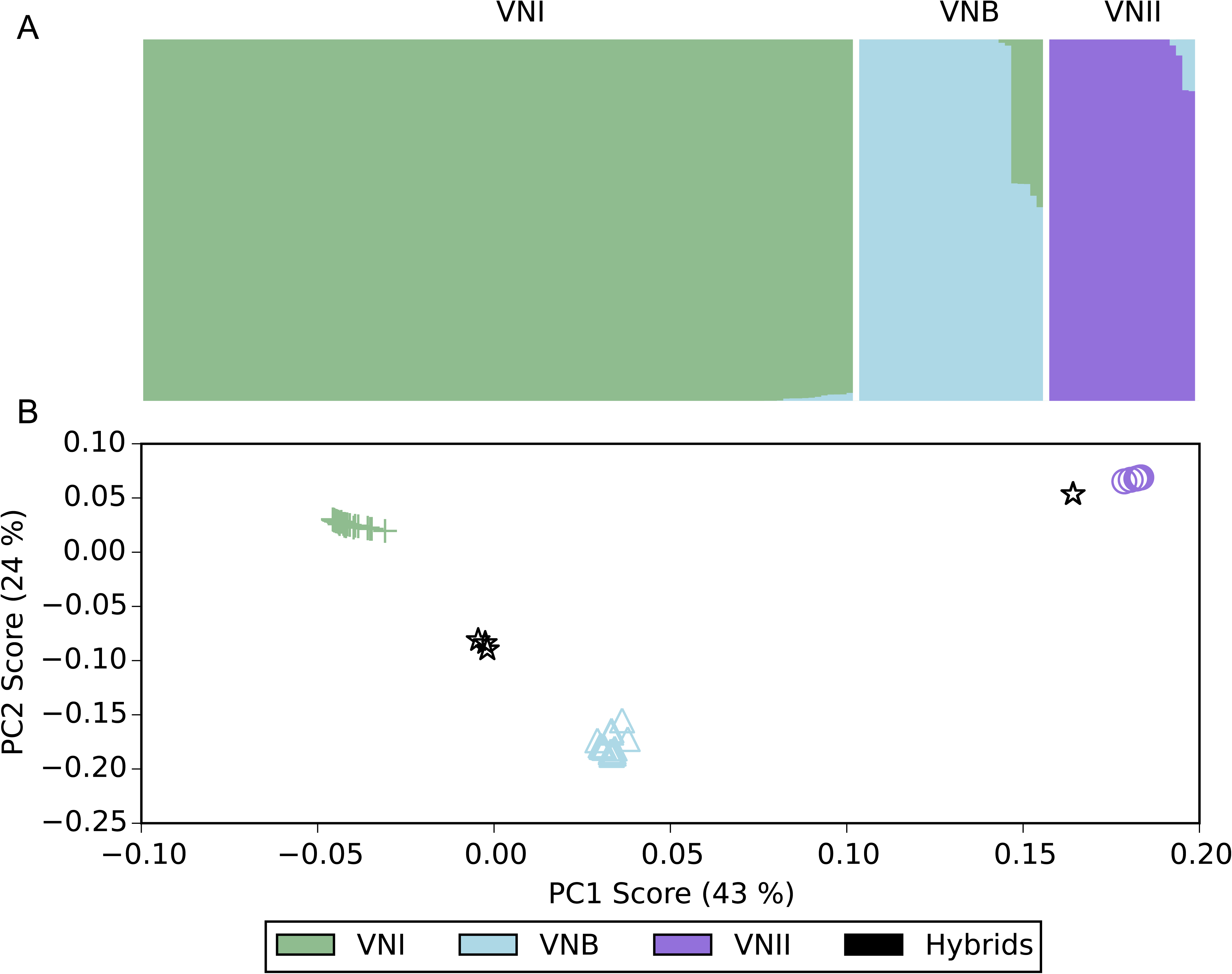
Ancestry characterization of three major groups highlights hybrid isolates. A. The fraction of ancestry (k=3) estimated by STRUCTURE is shown within a column for each isolate. B. Principal components analysis separates the 3 major lineages, with the hybrid isolates showing a mix of VNB ancestry with either VNI or VNII.

Next, we identified the ancestry contribution of each isolate using STRUCTURE with three population subdivisions. This confirmed that most isolates have a single dominant ancestry assigned to the VNI, VNB, and VNII lineages. In addition, the isolates with intermediate positions indicated by PCA were found to have mixed ancestry contributions by STRUCTURE. SNP sites for the VNI-VNB hybrids contain an average of 40.8% VNI ancestry and 59.2% VNB ancestry whereas the VNII-VNB hybrids have 85.8% VNII and 14.2% VNB ancestry (**Table S2**). The similar fraction of ancestry in the VNI-VNB hybrids suggests they could be recent mixtures of the two lineages, whereas the VNII-VNB hybrids may be more ancient mixtures with additional crosses to VNII isolates biasing the final ratio of parental SNPs.

### Evidence of recent meiotic exchange generating haploid hybrids

To examine the degree of intermixing of ancestry for these hybrid genotypes across the genome, we identified the most likely ancestry for each SNP site using the site-by-site mode in STRUCTURE. Selecting positions where the ancestry assignment was most confident (0.9 or greater, Methods), we examined the distribution of these sites by ancestry across the fourteen chromosomes (Figure 3). Each of the three VNI-VNB hybrids displayed different patterns of large regions corresponding to a single ancestry. For example, chromosome 1 has three large blocks of different ancestry in Bt125, four in Bt131, and two in Ftc260-1 (Figure 3A-C). While all chromosomes contained regions of both VNI and VNB ancestry groups in Bt125 and Ftc260-1, two chromosomes of Bt131, chromosome 6 and 9, have only large regions of VNB ancestry. By contrast, CCTP51, which contains a lower fraction of the second ancestry (VNB), appears more highly intermixed with smaller ancestry blocks (Figure 3D). Notably, three of the four unique genotypes (Bt131, CCTP51, and Ftc260-1) contain the rare *MAT***a** locus; in all *MAT***a** isolates, the mating type locus region is of VNB ancestry, whereas the mating locus region in the *MAT*α isolate (Bt125) is of VNI ancestry (Methods). Overall these patterns suggest a recent hybridization of VNI and VNB isolates, with recombination during meiosis generating chromosome-wide intermixing resulting in distinct parental haplotype blocks. In Bt125, a 205 kb region of scaffold 6 is present at nearly twice (1.92 fold) the average depth. Otherwise this isolate, and the other six hybrid isolates, was found to contain even levels of ploidy across the 14 chromosomes based on read depth.

**Figure 3.**
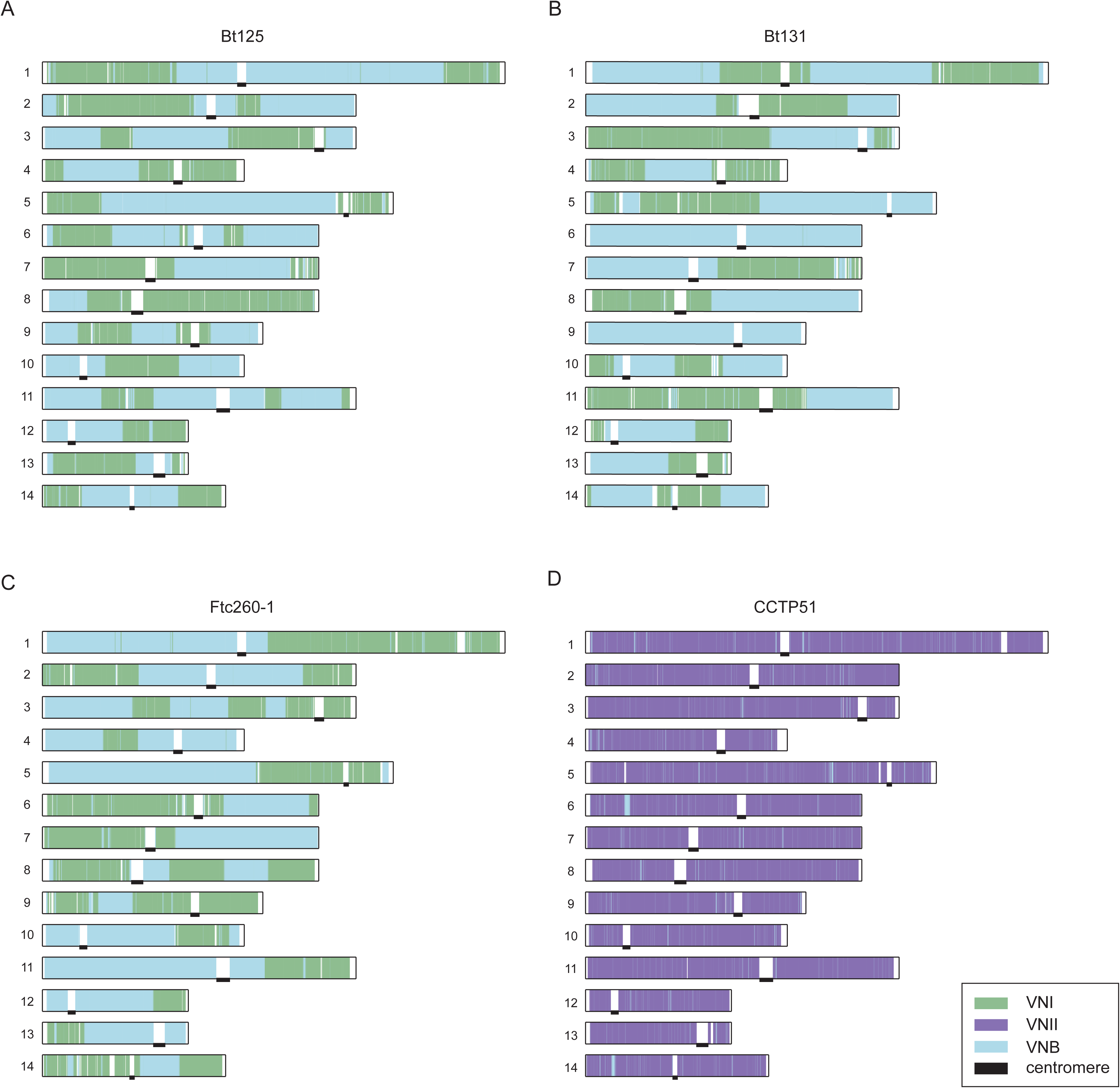
Large blocks of ancestry suggest recent recombination between lineages. For each of the four isolates depicted (A-D), the STRUCTURE assigned ancestry for each site along each chromosome is depicted as a colored bar corresponding to VNI, VNII, and VNB ancestry. Locations of centromeres are marked with black bars.

For the three VNI-VNB hybrids showing large ancestry blocks, we also utilized the site ancestry predictions to finely map the genotypes within each population. Given the roughly equal contribution of the two ancestry sites and the large block size for each in these genomes, we hypothesized that these hybrids could have resulted from recent mating of one genotype of each lineage, which we could reconstruct using separate phylogenies of each site class. For each genotype, sites mapped to either the VNI or VNB ancestry were selected and a separate phylogeny constructed for each of these two sets of sites. For VNI ancestry sites, these isolates had very different genotypes, with Ftc260-1 most closely related to a diverse set of African isolates in VNI, whereas both Bt125 and Bt131 are more closely related to highly clonal clades of VNI isolates (**Figure S2A,C,E**). Similarly for a separate phylogenetic analysis of VNB ancestry sites, Bt125 and Bt131 were placed within one of the two major subclades of VNB while Ftc260-1 was placed in the other (**Figure S2B,D,F**). This supports that these three hybrids originated from very different genotypes of VNI and VNB parental isolates.

### Diploid isolates and genome plasticity

As noted above, a total of 24 sequenced isolates displayed heterozygous SNP positions across the genome. Four of these isolates had higher rates of polymorphism overall and appear to be hybrids within or between VN lineages (Bt66, Cng9, PMHc.1045.ENR.STOR, and 102-14) (**Figure S3**). Each of these isolates contain two copies of the *MAT*α mating type locus, suggesting they arose from same sex mating of two *MAT*α parental isolates. In addition, 11 serotype A diploids showed very low rates of heterogyzosity (**Figure S3**), consistent with AFLP and MLST-based evidence that they arose from endoreduplication or self-mating (Lin et al., 2009). The remaining isolates include eight serotypeA/serotypeD diploids, of which seven contain both *MAT***a** and *MAT*α mating types and one is homozygous for the *MAT***a** locus, and one serotype A/*Cryptococcus gattii* hybrid containing two copies of *MAT*α.

All types of diploid isolates in our set, including A/A diploids, exhibit regions of loss of heterozygosity (LOH) in the genome, where alleles of only one parental isolate are present. Three of the A/A diploids (Bt66, Cng9, and 102-14) are heterozygous throughout nearly all of the genome; Cng9 exhibited only a small LOH region at the start of chromosome 2, which also has haploid levels of genome coverage. Isolate PMHc1045 by contrast has large LOH regions on six scaffolds, including a 1.1 Mb region of chromosome 6 (**Figure S3**). Some of these regions of LOH in PMHc1045 are linked to aneuploid chromosome segments, including a region of chromosome 12 reduced to haploid levels and or triploid levels of the region adjacent to a LOH on chromosome 6. All LOH regions are telomere-linked, reminiscent of what has previously been reported across diverse isolates of *Candida albicans* (Hirakawa et al., 2015).

We next inferred the ancestry of the two parental isolates contributing to the A/A hybrids by examining the frequency of SNP alleles that are highly predictive for VNI, VNII, or VNB (Methods). Three of the isolates (Cng9, PMHc1045, and 102-14) have similar frequencies of such VNII and VNB alleles, whereas Bt66 is comprised of VNI and VNB predictive alleles (**Table S3**). Comparing Cng9 and PMHc1045 directly, 89.2% of variant sites are identical; this fraction increases to 97.3% when LOH regions are excluded and a similar fraction of sites are shared with 102-14. By comparison, both isolates share 23% of variant positions with Bt66 when LOH regions are excluded from both. Notably, LOH has resulted in a mixing of genotypes; examining predictive alleles for each of the eight LOH regions of PMHc1045 revealed regions corresponding to each lineage. Two regions encompassing 1.4% of the genome share the highest fraction of private alleles with other VNB isolates whereas the remaining six regions encompassing 10.2% of the genome share most private alleles with other VNII isolates. Thus, LOH has led to large differences between otherwise highly similar Cng9 and PMHc1045 isolates and resulted in blended ancestry by converting regions to each of the two parents in PMHc1045.

The eight AD hybrids also showed evidence of even more extreme aneuploidy and LOH related to loss of one of the two parental chromosomes. All isolates displayed evidence of aneuploidy, by examining read coverage across both the H99 serotype A and JEC21 serotype D reference genomes (**Figure S4**). While some isolates have retained chromosomes of both A and D origin, others have lost a chromosome from one parent and duplicated the corresponding chromosome of the other (Figure 4, **Figure S4**). For example in RCT14, two copies of chromosome 1 are present but both have serotype A origin; similarly in IFNR21, both copies of chromosome 10 have serotype D origin. Both of these isolates display additional aneuploidies, with 3 copies of some chromosomes. Notably, CCTP50 appears mostly triploid, with either 2:1 or 1:2 ratios of the A:D ratio for each chromosome (Figure 4); this pattern is also observed in IFN26 (**Figure S4**). In IFN-R26, loss of chromosome 4 in JEC21, balanced by gain of chromosome 5 in H99 (**Figure S4**), has resulted in a *MAT***a**/*MAT***a** genotype. While the mating type of the original JEC21 parent can not be determined, this suggests that generation of *MAT***a**/*MAT***a** diploids can occur via chromosome loss and duplication. All other isolates are *MAT***a**/*MAT*α, suggesting that they originated from opposite sex mating. While diploid AD hybrids have been isolated from both environmental or clinical sources (Litvintseva et al., 2006), all eight AD hybrids in our set are of clinical origin.

**Figure 4.**
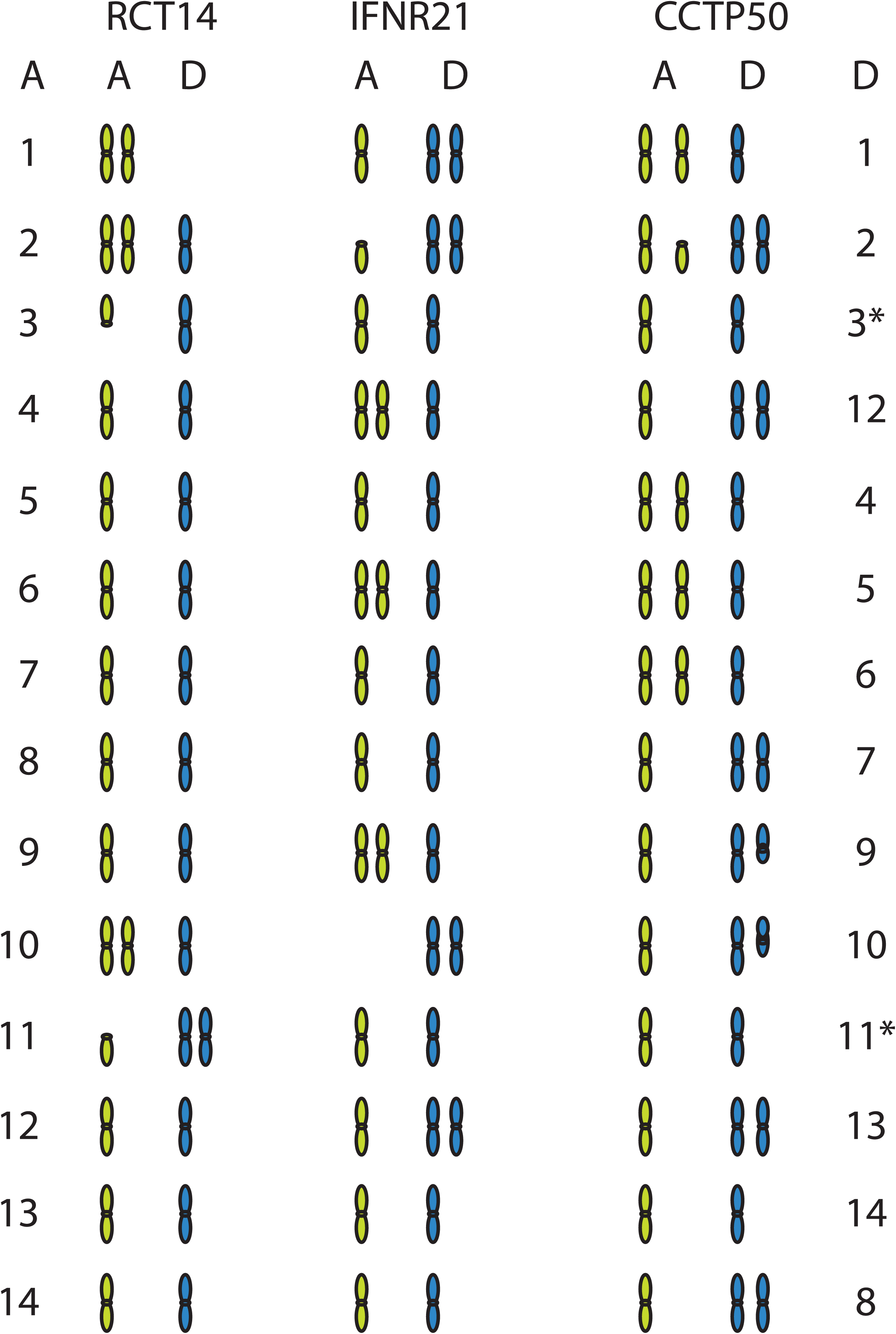
Chromosome ancestry and ploidy variation of AD hybrids. For three AD hybrid isolates (RCT14, IFNR21, and CCTP50), the contribution and copy number of A (green) and D (blue) ancestry chromosomal regions was measured by aligning all sequence reads to a combined AD reference (H99, left and JEC21, right). The copy number of each chromosome is depicted, with either full or partial chromosomal regions shown; see Figure S4 for detailed coverage plots for all AD hybrid isolates.

To examine the diversity of these AD hybrids, SNPs were identified by comparison to a combined A (H99) and D (JEC21) genome reference. Phylogenetic analysis of A and D genome SNPs revealed that both the A and D copies of each hybrid are closely related for these isolates (**Figure S5**). On average, the A genomes differ by 6,108 SNP positions and the D genomes by 3,935 SNP positions. The A genomes are from the VNB lineage, most closely related to Bt206 in our analysis (**Figure S5**). The low diversity of both the A and D genomes between isolates suggests that this set of 8 AD hybrids may have originated from a single hybrid isolate or from a set of closely related A and D parental isolates.

### Chromosomal copy number variation

On a smaller scale than whole-genome hybridization, chromosomal copy number variants appear to be common in *C. neoformans* and may be an adaptive mechanism for virulence (Rhodes et al., 2017). In the set of 164 primarily haploid isolates, 25 exhibited whole or partial chromosomal aneuploidies (**Figure S6**). In 13 of the 25 isolates, an entire chromosome or region thereof showed a doubling of sequencing coverage, consistent with a diploid chromosome in an otherwise haploid isolate. The remaining 12 isolates show a 50% gain in coverage better explained by a diploid isolate with a triploid chromosome or region. These likely diploid isolates do not display heterozygous base calls, suggesting a recent endoreduplication of the genome and associated aneuploidy of additional chromosomes.

Aneuploidies of particular chromosomes may provide a specific biological advantage or alternatively be better tolerated. In general, the smallest chromosomes (12 and 13) are the most frequently observed to exhibit aneuploidy (**Figure S6**). Several isolates have an increased copy number of chromosome 1; amplification of the lanosterol-14-α-demethylase *ERG11* and the major efflux transporter *AFR1* located on chromosome 1 can confer resistance to azole drugs (Sionov et al., 2010). Of the four isolates that contain chromosome 1 aneuploidies, either *ERG11* (CCTP34) or *AFR1* (IFN-R11 and RCT6) or both genes (CCTP9) are present at elevated copy number. The elevated copy number of *AFR1* appears correlated with increased drug resistance; both CCTP9 and RCT6 displayed fluconazole MIC values of 256 ug/ml, whereas CCTP34 appeared more susceptible at an MIC of 8 ug/ml (Methods). Notably, all of the isolates with chromosome 1 aneuploidies are of clinical origin, as are 24 of all 25 isolates with detected aneuploidies (**Figure S6, Table S1**). Of the seven isolates with hybrid ancestry, only Bt125 included a small region of chromosome 6 at higher copy number; otherwise this and the other hybrid isolates appeared to be haploid. Across the diploid and haploid isolates, we detected aneuploidies affecting all chromosomes (**Figures S3, S4, and S6**).

### Conservation of gene content and structure across lineages

To examine the extent of gene content variation across the three major lineages of *C. neoformans* var. *grubii*, we assembled and annotated genomes of 39 representative isolates (Methods). Previously a high quality reference genome was produced for the H99 VNI isolate (Janbon et al., 2014); our data set includes new annotated assemblies for 9 diverse VNB isolates, 27 VNI isolates, and three VNII isolates (**Table S4**). The gene sets across all 40 assemblies (including H99) were compared to each other and to those of four *C. gattii* (representing VGI, VGII, VGIII, and VGIV) and one *C. neoformans* var. *neoformans* (serotype D) reference genomes (Methods) in order to evaluate gene conservation. Based on orthologs identified across these genomes (Methods), an average of 4,970 genes are conserved across all 45 compared *Cryptococcus* gene sets; within serotype A, an average of 5,950 genes are conserved in all 40 genomes (**Figure S7**). A phylogeny inferred from 4,616 single copy genes supports VNII in an ancestral position relative to the more recently diverging VNI and VNB (**Figure S7**).

Gene content is highly conserved across *C. neoformans* var. *grubii* with few examples of genes specific to the separate lineages (**Supplementary Note**). Based on ortholog profiling, a total of 11 genes are specific to VNI, three specific to VNB, and 25 specific to VNII (**Table S5**). These include two clusters of genes specific to VNI or VNII located within otherwise syntenic regions of the genome (Figure 5). The cluster of five genes unique to VNI genomes include a predicted haloacid dehydrogenase, an amidohydrolase and an allantoate permease, which could be involved in uptake of uric acid products. The cluster of six genes unique to the VNII genomes includes a predicted transcription factor, amino acid transporter, hydrolase, dihydropyrimidinase, and oxygenase superfamily protein. While both clusters are also missing from the JEC21 *C. neoformans* var *neoformans* genome, the more distantly related *C. gattii* genomes contain syntenic orthologs of all of the VNII-specific cluster genes and between 1 and 3 non-syntenic orthologs of the VNI-specific cluster. These patterns suggest gene loss and perhaps lateral transfer in some species and lineages account for these differences. There was little other evidence of lineage-specific gene loss; orthologs missing in only one lineage included only hypothetical proteins. In addition, we further searched for genes with loss-of-function mutations in all members of each lineage using SNP data, to find genes that may be disrupted but still predicted in the assemblies. However, we found no convincing evidence of disrupted genes with known functions in any of the three lineages (**Supplementary Note**).

**Figure 5.**
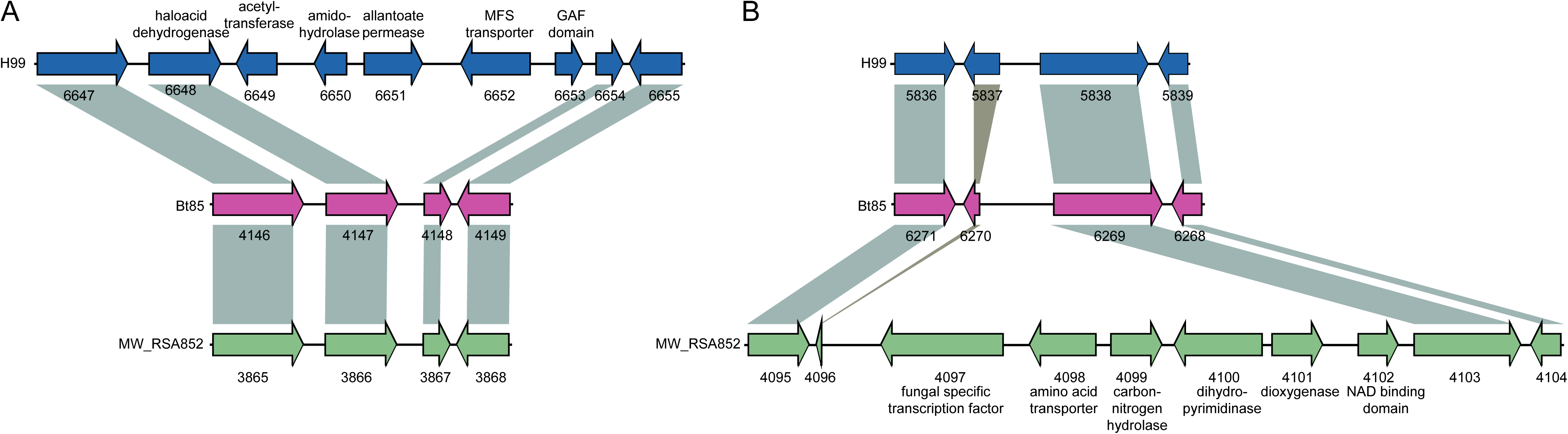
Lineage-specific gene clusters. Two large-lineage specific clusters were detected in the VNI genomes or VNII genomes; these are depicted using a representative genome from each lineage. A. Insertion of CNAG_06649 to CNAG_06653 in H99 (blue, VNI); syntenic genes in Bt85 (pink, VNB) and MW_RSA852 (green, VNII) are connected with grey bars. B. Insertion of C358_04097 to C358_04102 in MW_RSA852.

Given the high level of gene conservation between lineages, we sought to identify rapidly evolving genes that might be involved in phenotypic differences between *C. neoformans* lineages. For each gene, we built a consensus sequence for each lineage and then calculated pairwise *d*_N_ and *d*_S_ of these fixed sites. As *d*_S_ was uniformly low throughout the dataset due to limited genetic diversity, we identified differences in *d*_N_, which measures both the mutation rate and selection. The top 10 annotated genes with the largest *d*_N_ for each pairwise comparison are shown in Table 2, and the three comparisons in total include 18 unique genes. The set is dominated by transcription factors (*GLN3*, *PDR802*, *SXI1α*, *YOX101*, and *ZNF2*) and transferases (*ATG2602*, *CDC43*, *GPI18*, *HOC1* and *RAM1*), many of which have already been implicated in virulence (Esher et al., 2016; Jung et al., 2015; Lee et al., 2015; Selvig et al., 2013; Wang et al., 2012) or resistance to oxidative stress (Jung et al., 2015). In particular, *CDC43* and *RAM1* are both rapidly evolving; these genes represent the two major independent methods of prenylation, key in proper subcellular localization of many proteins, often to the membrane (Esher et al., 2016; Selvig et al., 2013). Other rapidly evolving genes include β-glucan synthase *KRE63,* superoxide dismutase *SOD1*, and mating regulator *SXI1α*, the latter of which is highly divergent between VNII and both VNI and VNB, and could play a role in reproductive isolation of the VNII lineage.

**Table 2.** Rapidly evolving genes in the three lineages of *C. neoformans* var. *grubii*. Consensus sequences were built for each lineage, and *d*_N_ and *d*_S_ were calculated for each lineage pair. As *d*_S_ was uniformly low throughout the dataset due to limited genetic diversity, for each pair of lineages we identified the 10 genes with assigned names (Inglis et al., 2014) with the highest *d*_N_, which measures both the mutation rate and selection.

| Comparison | $d_N$ | Locus | Gene | Annotation |
| --- | --- | --- | --- | --- |
| VNI vs VNB | 0.0181 | CNAG_01841 | *GLN3* | transcription factor, deletion sensitive to organic peroxides (Jung et al., 2015) |
|  | 0.0155 | CNAG_03894 | *PDR802* | transcription factor, deletion with reduced virulence (Jung et al., 2015) |
|  | 0.0095 | CNAG_03213 | *UVE1* | UV damage endonuclease |
|  | 0.0092 | CNAG_02756 | *CDC43* | geranylgeranyltransferase-I, essential for virulence (Selvig et al., 2013) |
|  | 0.0090 | CNAG_06655 | *GPI18* | GPI-anchor transamidase |
|  | 0.0089 | CNAG_01908 | *HEM4* | uroporphyrinogen-III synthase |
|  | 0.0085 | CNAG_03133 | *ATG2602* | UDP-glucose sterol transferase |
|  | 0.0084 | CNAG_03617 | *CLP1* | clampless protein 1 |
| | 0.0076 | CNAG_05740 | *RAM1* | farnesyltransferase β-subunit, essential for virulence (Esher et al., 2016) |
|  | 0.0068 | CNAG_03637 | *YKU80* | Double strand break repair factor and silencing regulator, deletion has reduced virulence (Liu et al., 2008) |
| VNI vs VNII | 0.0610 | CNAG_05836 | *HOC1* | $\alpha$ 1,6-mannosyltransferase (Lee et al., 2015) |
|  | 0.0408 | CNAG_05838 | *RGD1* | Rho GTPase activating protein, deletion has increased virulence (Liu et al., 2008) |
| | 0.0214 | CNAG_06031 | *KRE63* | β-glucan synthase, involved in capsule and cell wall formation, deletion has decreased virulence (Gilbert et al., 2010) |
| | 0.0149 | CNAG_06814 | *SXI1α* | $\alpha$ cell type transcription factor, required for mating (Hull et al., 2002) |
|  | 0.0142 | CNAG_01841 | *GLN3* | see above |
|  | 0.0135 | CNAG_03229 | *YOX101* | transcription factor, deletion sensitive to organic peroxides (Jung et al., 2015) |
|  | 0.0127 | CNAG_03398 | *ZIP2* | zinc ion transporter |
|  | 0.0113 | CNAG_03133 | *ATG2602* | see above |
|  | 0.0110 | CNAG_03366 | *ZNF2* | transcription factor, overexpression results in reduced virulence (Wang et al., 2012) |
|  | 0.0104 | CNAG_01019 | *SOD1* | superoxide dismutase |
| VNB vs VNII | 0.0617 | CNAG_05836 | *HOC1* | see above |
|  | 0.0402 | CNAG_05838 | *RGD1* | see above |
|  | 0.0171 | CNAG_06031 | *KRE63* | see above |
|  | 0.0128 | CNAG_03366 | *ZNF2* | see above |
|  | 0.0122 | CNAG_06814 | *SXI1α* | see above |
|  | 0.0114 | CNAG_03213 | *UVE1* | see above |
|  | 0.0104 | CNAG_01019 | *SOD1* | see above |
|  | 0.0104 | CNAG_03398 | *ZIP2* | see above |
|  | 0.0102 | CNAG_01841 | *GLN3* | see above |
|  | 0.0102 | CNAG_02756 | *CDC43* | see above |

### Population measures and biogeography

Strikingly, recently identified VNB genotypes from South America are placed in the phylogeny as basally branching clades for each VNB subgroup, which otherwise consist of genotypes from Africa (Figure 1). All of the six South American VNB isolates contain the *MAT*α genotype. By contrast, both VNI and VNII consist of more closely related though more geographically diverse sets of isolates; one large clonal group is found in VNII, whereas several are observed for VNI, which is oversampled owing to its higher prevalence in patients and environments worldwide. Overall, VNB showed the highest average pairwise diversity (pi=0.00736), nearly four times the level in VNI (pi=0.00200), with the lowest value for VNII (pi=0.00105) (Table 3). Genetic diversity within the VNB lineage was similar between the South America and African isolates (pi=0.00727 and 0.00736, respectively). However, genetic diversity of VNI isolates in India was lower than VNI isolates in Africa (pi=0.00146 and 0.00337). VNB also contained the largest fraction of private alleles compared to VNI and VNII, reflecting the higher variation within VNB (Table 4). By contrast, VNI and VNII had the highest number of fixed differences, reflecting the long branches leading to these clades. The average divergence (dXY) between lineages ranges is 0.012 comparing isolates from VNI and VNB and 0.015 for comparison of either to VNII (Table 4), highlighting the low nucleotide divergence between the lineages. VNI and VNII were the most differentiated of the three lineages as shown by pairwise whole genome fixation indexes (*F*_st_) (Weir and Cockerham 1984). The highest average chromosome *F*_st_ value is 0.874 between VNI and VNII isolates, while the average chromosome *F*_st_ values of VNI-VNB and VNB-VNII are 0.595 and 0.707, respectively (Table 4).

**Table 3.** Comparison of VN group statistics

| Populations | Isolates<br>(#) | Segregating<br>sites | Pi | Tajima's D |
| --- | --- | --- | --- | --- |
| VNI | 111 | 190,716 | 0.00200 | -0.107179 |
| VNII | 23 | 337,990 | 0.00105 | -1.005950 |
| VNB | 25 | 613,991 | 0.00736 | -0.232596 |

**Table 4.** Pairwise comparison of VN group statistics

| Comparisons | Fixed | Shared | Private_A | Private_B | dXY | Fst |
| --- | --- | --- | --- | --- | --- | --- |
| VNB vs VNI | 54,719 | 52,536 | 446,566 | 102,817 | IvB: 0.0119 | IvB: 0.595 |
| VNB vs VNII | 118,329 | 68,211 | 405,406 | 78,444 | BvII: 0.0154 | BvII: 0.707 |
| VNI vs VNII | 188,590 | 38,501 | 116,845 | 83,802 | IvII: 0.0152 | IvII: 0.874 |

To further examine the evolutionary history of the novel South American VNB isolates, we subdivided VNB into four subclades (VNBI-South America, VNBI-Africa, VNBII-South America, and VNBII-Africa) and calculated alleles unique to each subclade and shared across VNB groups or geography (Methods). These subclades represent all combinations of the two previously identified VNB groups (VNBI and VNBII) and the two geographies (South America and Africa). One South American VNB isolate (V53), nested deeply within African isolates on the phylogeny, was excluded from the analysis. All four of the subclades contained more unique alleles than were shared across either VNB group or geography (Figure 7), suggesting both a high level of genetic diversity within each subclade and some degree of reproductive isolation between each subclade. Furthermore, there was greater number of unique alleles shared within the VNB groups from different geographic regions than were shared across VNB groups within the same geographic region (Figure 7). This geographically and phylogenetically segregated diversity suggests that one or more ancient migration events occurred between South America and Africa during the diversification of VNB, followed by geographic isolation. In contrast, the VNI and VNII lineages showed a pattern consistent with more rapid current migration, where isolates from different geographic regions in many cases differed by fewer than 200 SNPs.

We next evaluated whether VNI and VNB showed a signal of genetic isolation by distance using the Mantel test. In both VNI and VNB, genetic distance was significantly positively correlated with geographic distance (p = 0.0001 and p = 0.042, respectively). When VNB was separated into VNBI and VNBII, each lineage showed an even stronger signal (p = 0.0051 and p = 0.0009, respectively), suggesting much of the correlation seen within VNB is representative of isolation within each subclade. Therefore, despite VNB showing signals of more ancient migration while VNI shows signals of recent migration, both demonstrate genetic substructure according to geography.

### Recombination between and within lineages

The basal branching of Brazilian VNB isolates revealed in the phylogenetic analysis suggested that South America could be a global center of *C. neoformans* var. *grubii* diversity. To further investigate this hypothesis, and to explore recombination in the context of population structure, we implemented the chromosome painting approach of fineStructure (Lawson et al., 2012), which identifies shared genomic regions between individuals and thereby ancestral relationships among individuals and populations. Our linked co-ancestry model found the highest level of sharing among VNB isolates; in addition, there is evidence of strong haplotype donation from South American VNB isolates (V2, V31, and V87) to all other lineages and continents, suggestive of ancestral recombination (Figure 6).

**Figure 6.**
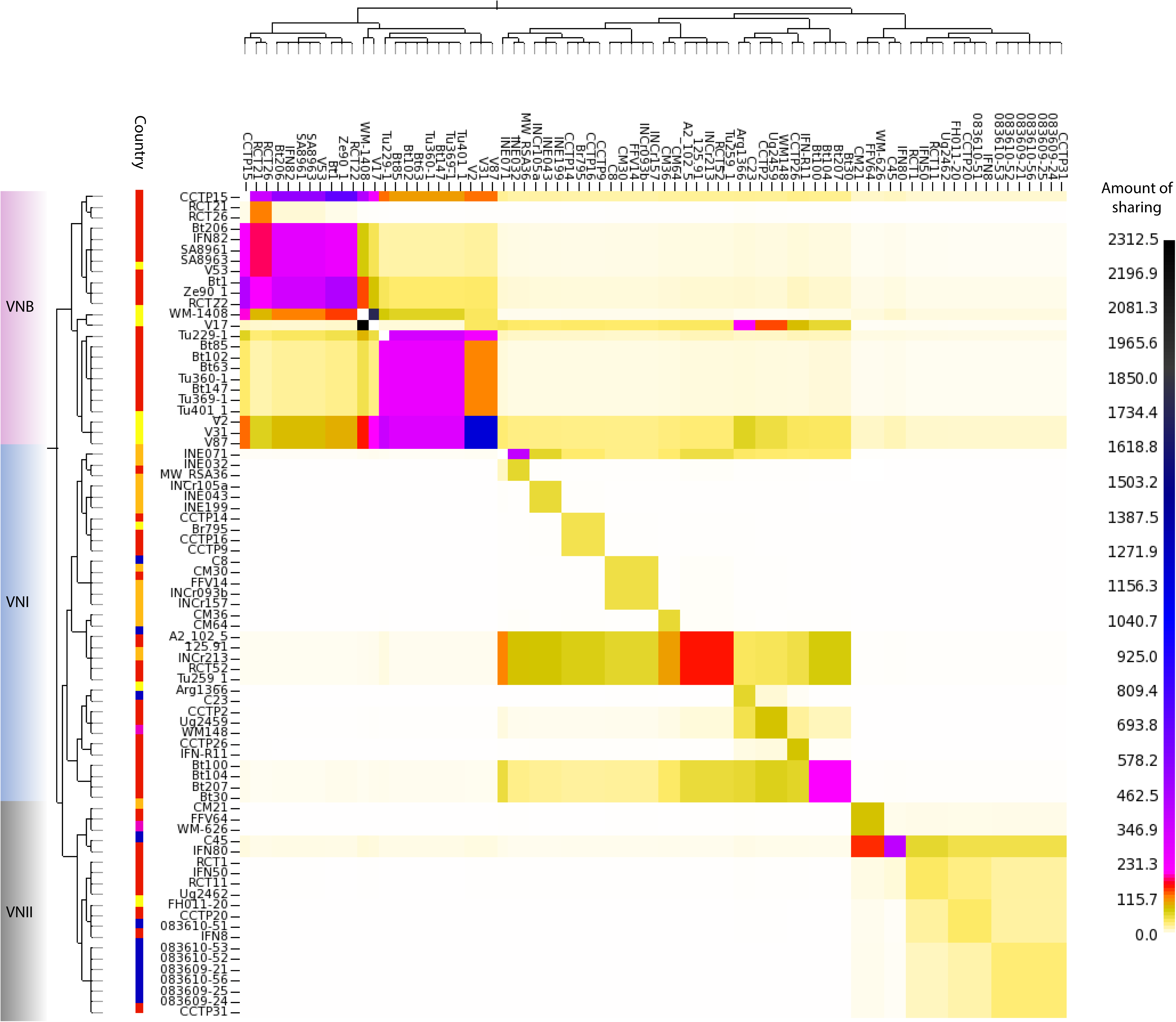
Genome-sharing analysis of *Cryptococcus neoformans* var. *grubii* using fineStructure was performed on a SNP matrix using a representative of each clonal population within the VNI lineage. These genomes were reduced to a pairwise similarity matrix, which facilitates the identification of population structure based on haplotype sharing within regions of the genome. The *x-*axis represents the “donor” of genomic regions, while the *y*-axis represents the recipient of shared genomic regions. The scale bar represents the amount of genomic sharing, with black representing the largest amount of sharing of genetic material, and white representing the least amount of shared genetic material (no sharing). The geographical site of isolation is illustrated with coloured boxes as in Figure 1, and lineages are also shown.

**Figure 7.**
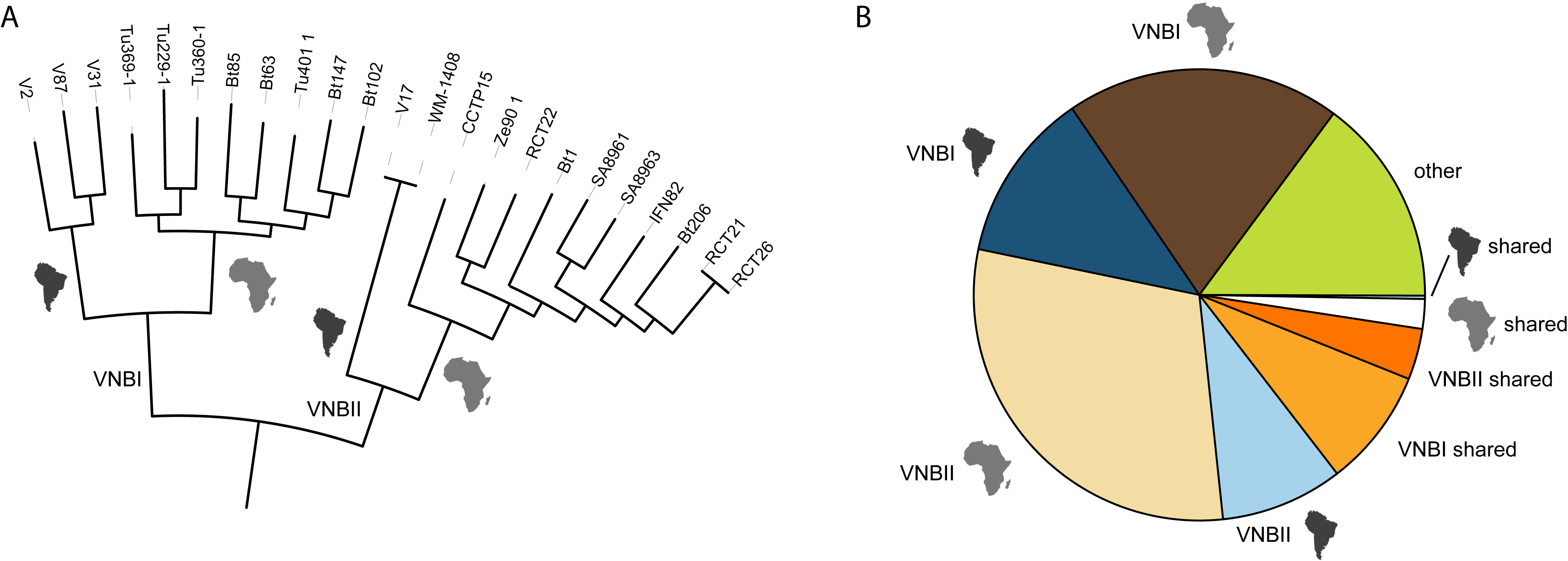
VNB alleles in population subdivisions and across geography. A. Phylogeny of VNB lineage showing major subdivisions (VNB-I and VNB-II) and inferred ancestral geography (South America or Africa, depicted as continent shapes). B. Classification of all 445,193 private VNB alleles (present in at least 1 VNB isolate and no VNI or VNII isolates) by subdivisions and geography. Most VNB alleles are specific for the each VNB subdivision and for the geographic subdivisions within each group. More alleles are shared between geographic locations in the same subdivision (VNBI or VNBII) than are shared within geographic locations across subdivisions.

Independent confirmation of ancestry using STRUCTURE confirmed that V87 includes primarily VNB ancestry with ∼1% VNI alleles (**Table S7**). Interrogating the chunk counts, which are lengths of DNA shared by a donor to other individuals, and lengths produced by fineStructure revealed that the haplotype chunks donated by these ‘ancestral’ isolates were substantially higher than seen for other isolates, with other African VNB isolates receiving significant chunks and lengths (Bt102, Bt63, Bt85, Tu229-1, Tu360-1, Tu369-1, and Tu401-1) from the South American VNB isolates. Isolate V53 donated less strongly than these three isolates to all lineages. Other South American VNB isolates (WM 1408 and V17) donated strongly to specific lineages: WM 1408 to VNII and VNB, whilst V17 donated to VNI and VNB. However, these findings for WM 1408 and V17 were not corroborated using STRUCTURE. Despite their allocation to separate VNB subpopulations, V2 and V17 (VNB-I and VNB-II respectively) donate the most genetic material (when interrogating the chunk counts) to VNI isolates in Africa, India, and Thailand.

Within the VNI lineage, fineStructure analysis identified a subset of isolates with a high frequency of haplotype sharing (Figure 8). Notably, a group of African (Tu259-1, 125.91, RCT52, Bt100, Bt207 and Bt30) and Indian (INCr213 and INE071) isolates show strong haplotype donation with many other VNI isolates, suggestive of ancestral recombination events. These isolates are dispersed over four subpopulations within the VNI lineage. Though the geographical distance between these populations should preclude frequent intermixing, these isolates from Africa and India may include a higher fraction of ancestral alleles, leading to a lack of phylogeographic structure among these highly geographically distinct populations.

**Figure 8.**
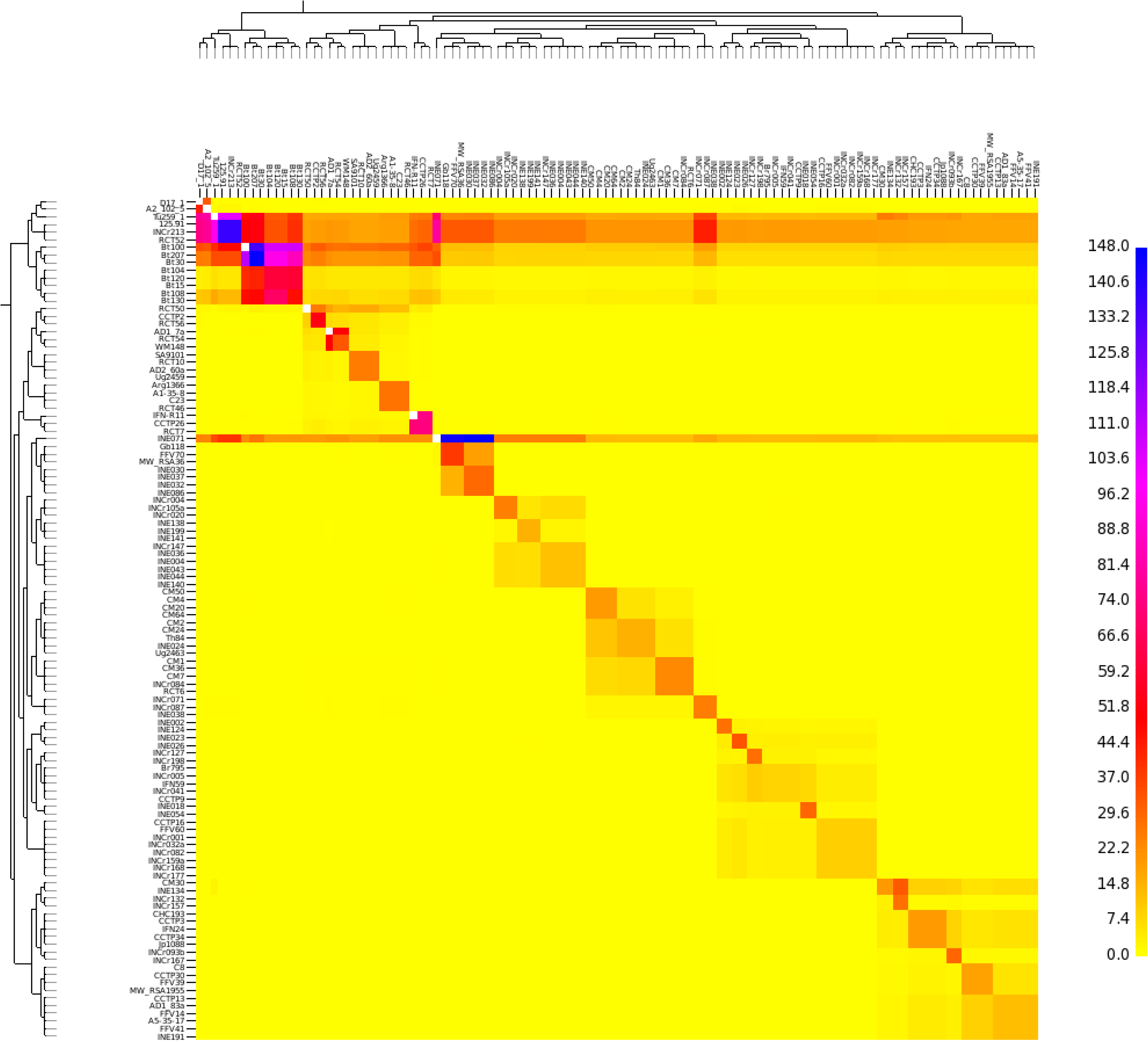
Genome-sharing analysis of the *Cng* VNI lineage using fineStructure on a SNP matrix of 111 genomes. The *x-*axis represents the “donor” of genomic regions, whilst the *y*-axis represents the recipient of shared genomic regions. The scale bar represents the amount of genomic sharing, with blue representing the largest amount of sharing of genetic material, and yellow representing the least amount of shared genetic material (no sharing).

Finding that ancestral recombination in the VNB lineage contributed to VNI lineage diversity suggested that there will be a signature of admixture linkage disequilibrium (LD) in these two populations. Linkage disequilibrium differs between lineages (**Figure S8**), with VNII LD decaying slowly with physical distance, and manifesting an LD50 (where linkage disequilibrium has decayed to half its maximum value) at >150 kb. However, this value may reflect the highly clonal nature and relatively small number of sequenced VNII isolates. LD decay is relatively slow for VNI with an LD50 of 4,500 bp, whereas LD decays more rapidly in the VNB lineage, with an LD50 of 1,500 bp. When separated into geographical origin of isolation (**Figure S8** (b)), LD50 for South American VNB appears greater (> 150 kbp) than that seen in African VNB (2,000 bp). The slower decay of LD in VNI and VNII relative to VNB may reflect a predominately asexual mode of reproduction owing to the rarity of the *MAT***a** idiomorph.

### Discussion

This population genomic analysis of *Cryptococcus neoformans* var. *grubii* has revealed new biogeographic relationships and highlighted a complex history of hybridization events between groups. Analysis of genome-wide variation of 188 geographically diverse isolates greatly increases the resolution of the VNI, VNII, and VNB phylogenetic groups and precisely measures the level of genetic differentiation between isolates within each group and across geographic scales. This data supports a much higher diversity of isolates in the VNB group compared to VNI and VNII isolates. Notably, we show that hybridization between these groups can result in genome mixing suggestive of recent and ongoing meiotic exchange. A recent taxonomic proposal to divide the *C. neoformans* and *C. gattii* species complexes into seven monophyletic species did not subdivide *C. neoformans* var *grubii* into separate species; although VNI, VNII, and VNB were strongly supported clades in a multilocus phylogeny, coalescent based approaches did not clearly support these three lineages as separate species (Hagen et al., 2015). Our genome-wide analysis has uncovered new biogeographic structure and ongoing hybridization between lineages of *C. neoformans var. grubii*, suggesting that further subdivision is not straightforward. In addition, such hybridization events may be a biological feature that extends across other lineages within the *C. neoformans* and *C. gattii* species complexes (Farrer et al., 2015; Hagen et al., 2015), prompting a need for wider investigation of the population genomic structure of this whole complex is needed to support formal changes in taxonomy (see perspective by Kwon-Chung’17).

The placement of isolates from Brazil at basal branching positions of the two VNB subclades phylogenetically separates the South American and African isolates within both the VNBI and VNBII groups. This finding, along with the presence of a large number of unique alleles in each of these four subclades and strong haplotype sharing seen with fineStructure analysis (Figure 7), suggests that there was ancient migration of the VNB group between Africa and South America following the initial divergence of VNBI and VNBII but prior to each group’s diversification. This finding appears consistent with a prior report of diverse isolates from Brazil in a new VNI genotype 1B (Oliveira et al., 2004). While the lack of a reliable molecular clock combined with substantial rates of recombination prevents dating the time of divergence between VNB from South America and Africa with confidence, this division clearly occurred after these continents split over 110 million years ago, and also after VNB itself subdivided into two lineages – VNBI and VNBII. As is the case with VNI, cross-Atlantic migration events may have also vectored VNB between these two continents. However, despite evidence for these migration events, the majority of VNI and VNII migrations were likely much more recent than is seen with VNB, with nearly clonal isolates of VNI and VNII found in disparate geographic regions. The presence of one South American VNB isolate (V53) nested within African isolates on the phylogeny does, however, suggest a limited number of more recent migration events may be occurring between the two regions even within VNB despite the large degree of reproductive isolation that we observed. Identification of additional South American VNB isolates is necessary to determine their diversity and relationship to isolates from African continental regions. Although the sequenced isolates all contain the *MAT*α genotype, our sample size was small and likely under-represents the true diversity of this lineage in South America and the ecological reservoirs that it occupies.

Given the propensity of *Cryptococcus neoformans* var. *grubii* VNI and VNII for having an environmental reservoir in bird excreta (unlike VNB which is principally associated with arboreal reservoirs (Litvintseva et al., 2011; Vanhove et al., 2017)), it has been proposed that pigeons globally dispersed *Cryptococcus neoformans* var. *grubii* from a genetically diverse population in southern Africa (Litvintseva et al., 2011), resulting in an expansion of the *Cng* VNI out of Africa. Litvintseva et al. (2011) hypothesized that this “out-of-Africa” model for the evolution of VNI explains the origin of the global VNI population. Other studies showing lower genetic diversity of VNI populations in Southeast Asia (Simwami et al., 2011) and in South America (Ferreira-Paim et al., 2017), further supporting an African origin of *Cryptococcus neoformans* var. *grubii*. An alternative explanation for the higher diversity of African VNI could be that this lineage originated elsewhere and became more diverse in this continent by mating with the ‘native’ VNB population or due to other factors. Our analysis did not find a large subset of VNB alleles within the African VNI isolates based on ancestry analysis. In addition, we found one VNI subclade composed mostly of African isolates that appears to be recombining at higher frequency than other VNI groups. The phylogenetic intermixing of isolates from India and Africa strongly support the hypothesis that there is long-range dispersal and ancient recombination in environmental populations in India and Africa, indicative of multiple migratory events across time and into the present. Did VNI therefore evolve ‘Out of Africa’? Further sampling of environmental isolates from across South America as well as more diverse regions of Africa, as well as estimation of the mutation rate in *Cryptococcus neoformans* var. *grubii* to allow calibration of a molecular clock, is needed to further test this hypothesis.

While gene content is very similar across the *Cryptococcus neoformans* var. *grubii* lineages, we found examples of lineage specific genes including clusters unique to VNI or VNII. While this suggests that the *Cryptococcus neoformans* var. *grubii* gene inventory based on H99 (Janbon et al., 2014) is largely representative of all lineages, additional genes specific to VNII and VNB are important to consider in studies focusing on isolates of these lineages. Differences in gene expression may also differentiate the lineages, and it is important to note that these will include lineage-specific genes that may contribute to variation in clinical profiles and virulence that occur among lineages of *Cryptococcus neoformans* var. *grubii* (Beale et al., 2015). In addition, we found the most rapidly evolving genes between each of the lineages include transcription factors and transferases, suggesting phenotypic diversity may be generated through transcriptional reprogramming and protein modification rather than changes in gene content. The *SXI1* gene detected in comparisons of VNII with both VNI and VNB appears to be highly substituted in the VNII lineage; this sequence divergence of *SXI1* in VNII could contribute to differences in mating with this group. Truncated alleles of *SXI1* are frequently observed in the serotype D *MAT*α chromosome of AD hybrids and suggested to contribute to increased mating efficiency (Lin et al., 2007).

Our analysis revealed that hybrid isolates originate from multiple lineages, and resolved the parental genotypes. Prior analysis with MLST loci suggested that some isolates contain a mix of multiple genotypes (Chen et al., 2015; Litvintseva et al., 2003). However the sensitivity and precision of these methods is limited by the small number of loci examined, the use of genes involved in virulence that may be under different selective pressure, as well as incomplete lineage sorting in some groups. Analysis of genome-wide variation revealed that some isolates appear to be a recent mix of different ancestries, based on the detection of large blocks of sites with each ancestry; this could result from a small number of crossing over events for each chromosome during meiosis. Other isolates contain more highly intermixed ancestry across the genome and predominantly of a single ancestry; these may have occurred by more historical hybridization followed by subsequent mating within a single lineage group. The demonstration of genome mixing in hybrid isolates raises interesting questions about whether such fundamentally new assortments of the three lineages could generate genotypes with new phenotypes, which perhaps have a fitness or selective advantage.

Analysis of hybrids between serotypes A and D revealed a remarkable degree of genome reassortment. All of the 8 sequenced AD isolates show evidence of aneuploidy, affecting the copy number of 12 of 14 serotype A derived chromosomes and all 14 serotype D derived chromosomes. This is consistent with the high rate of AD isolate aneuploidy previously reported using flow analysis of DNA content (Lengeler et al., 2001) or comparative genome hybridization (Hu et al., 2008). For some chromosomes, only one parental genotype was detected in a subset of five isolates; this includes a loss of the serotype D copy of chromosome 1, as previously observed in analysis of three AD hybrid isolates (Hu et al., 2008). However, we further find that loss of heterozygosity (LOH) in some cases is due to partial copies of several chromosomes, suggesting that genomic instability in AD hybrids may result in chromosomal breakage. LOH was also observed for smaller regions in diploid AA hybrids. Similar LOH events are frequently observed in diploid fungi including *Candida albicans* (Hirakawa et al., 2015) and may contribute to the generation of genetic diversity in both species.

Aneuploidy was also commonly observed in the haploid *Cryptococcus neoformans* var. *grubii* isolates. Additional copies of regions of Chromosome 1 that include *AFR1* or *ERG11* are associated with drug resistance, though aneuploidies of additional chromosomes are also observed (Sionov et al., 2010). Although functional significance of aneuploidy of other chromosomes is less well understood, most of the isolates exhibiting aneuploidy are of clinical origin, suggesting increased copy of other genes may provide an advantage or that there is higher genome instability during infection. An isochromosome of the left arm of chromosome 12 that arose during infection has been reported (Ormerod et al., 2013) and chromosome 12 aneuploidy is widely seen in African patients with relapsed infections (Chen et al., 2017; Rhodes et al., 2017) although the specific role of this duplication is unclear. Our data suggests that there could be additional isochromosomes based on the detection of partial chromosomes using sequencing read depth; alternatively these regions could be represented in the genome as fusions with other chromosomes.

Previous studies of *Cryptococcus gattii* have pointed towards South America as a source of the diversity for the *C. gattii* VGII lineage (Engelthaler et al., 2014; Hagen et al., 2013). Given the shared evolutionary history of *C. gattii* and *Cryptococcus neoformans* var. *grubii* (Xu et al., 2000), South America could also represent a major diversity center of *Cryptococcus neoformans* var. *grubii*. Our data suggests that *Cryptococcus neoformans* var. *grubii* VNB isolates in both subgroups from South America have undergone ancestral recombination events, donating genetic material to all lineages across multiple geographical locations. Our study also provides clear evidence for recombination within the VNI and VNII lineages, where nearly all the isolates contain the *MAT*α mating type. This suggests that mating likely occurs between *MAT*α isolates, as is found in *C. neoformans* var. *neoformans* (Sun et al., 2014). Previous studies have hypothesized that *Cryptococcus neoformans* var. *grubii* can complete its sexual reproductive life cycle in environmental niches, such as plants (Xue et al., 2007) and pigeon guano (Nielsen et al., 2007; Vanhove et al., 2017). Our observations that all lineages of *Cryptococcus neoformans* var. *grubii* show the ability to widely disperse, to recombine, and to hybridize, illustrates that this pathogen has a high degree of evolutionary plasticity that is likely related to its success in infecting the immunosuppressed ‘human environment’, thereby causing a high burden of mortality worldwide (Armstrong-James et al., 2014).

### Methods

#### Isolate selection

A total of 188 *C. neoformans* var. *grubii* isolates were selected from previous studies, which include 146 clinical isolates, 36 environmental isolates, 4 animal isolates and 2 isolates of unknown isolation source; these isolates were collected from 14 different countries: Argentina, Australia, Botswana, Brazil, China, Cuba, France, India, Japan, South Africa, Tanzania, Thailand, Uganda and USA (**Table S1**). Most of the clinical isolates were isolated from the cerebrospinal fluid of patients. Eight of the 36 environmental isolates were isolated from pigeon guano, and most of the remaining isolates were collected from Mopane and other tree species.

#### Details of clinical trials and ethical review

French isolates were collected during the Crypto A/D study (Dromer et al., 2007). The study was approved by the local ethical committee and reported to the French Ministry of Health (registration # DGS970089). For clinical trials undertaken in South Africa (Bicanic et al., 2007, 2008; Jarvis et al., 2012; Loyse et al., 2012) and Thailand (Brouwer et al., 2004), ethical approval was obtained from the Wandsworth Research Ethics Committee covering St. George’s University of London. Local ethical approval was obtained from the University of Cape Town Research Ethics Committee in South Africa the ethical and scientific review subcommittee of the Thai Ministry of Public Health. Clinical isolates from India were collected during routine diagnostic service; local ethical approval was obtained from the Institutional Ethical Committee of Vallabhbhai Patel Chest Institute, University of Delhi, India.

#### Fluconazole sensitivity testing

Fluconazole MICs were determined for two isolates by the NHLS laboratory in Green Point, Cape Town using the E-test method (Biomerieux) (Bicanic et al., 2006).

#### DNA isolation and sequencing

Each yeast isolate was recovered from a freezer stock and purely cultured on an YPD or SD agar plate for 48-60 h. Next, a single colony was inoculated to another YPD plate and cultured for 24 h. Approximately 100 μl of yeast cells were used for DNA isolation using the MasterPure yeast DNA purification kit (Epicenter, Madison, WI) according to the manufacturer’s instructions. Alternatively, a single colony was inoculated into 6ml YPD broth supplemented with 0.5M NaCl and cultured for 40 hours at 37°C, prior to extraction using the MasterPure Yeast DNA purification kit (Epicentre) as previously described (Rhodes et al., 2017).

DNA was sequenced using Illumina technology; for each isolate, a small insert library was constructed and used to generate between 14 and 150 Million paired-end reads with 101bp per isolate, which results in 56 to 603 fold average coverage of reads aligned to the H99 genome. In addition, large insert libraries were constructed for 15 isolates (**Table S4**) and also used to generate 101bp paired-end reads. Isolates were sequenced at Imperial College London and the Broad Institute (**Table S1**).

#### Read alignment, variant detection, and ploidy analysis

Illumina reads were aligned to the *Cryptococcus neoformans* var. *grubii* reference genome H99 (Janbon et al., 2014) using the Burrows-Wheeler Aligner (BWA) 0.7.12 mem algorithm (Li, 2013) with default parameters. BAM files were sorted and indexed using Samtools (Li et al., 2009) version 1.2. Picard version 1.72 was used to identify duplicate reads and assign correct read groups to BAM files. BAM files were locally realigned around INDELs using GATK (McKenna et al., 2010) version 3.4-46 ‘RealignerTargetCreator’ and ‘IndelRealigner’.

SNPs and INDELs were called from all alignments using GATK version 3.4-46 ‘HaplotypeCaller’ in GVCF mode with ploidy = 1, and genotypeGVCFs was used to predict variants in each isolate. All VCFs were then combined and sites were filtered using variantFiltration with QD < 2.0, FS > 60.0, and MQ < 40.0. Individual genotypes were then filtered if the minimum genotype quality < 50, percent alternate allele < 0.8, or depth < 10.

In examining isolates with a high proportion of sites that were removed by these filters, inspection of the allele balance supported that these isolates were diploid. For heterozygous diploid isolates, haplotypeCaller was run in diploid mode. VariantFiltration was the same, with the added filter of ReadPosRankSum < −8.0. Then for individual genotype filtration there was no allele depth filter but otherwise was the same. The filters were kept as similar as possible to maximize combinability. For AD hybrids, a combined reference of H99 (Janbon et al., 2014) and JEC21 (Loftus et al., 2005) was used for alignment and SNP identification.

To examine variations in ploidy across the genome, the depth of bwa alignments at all positions was computed using Samtools mpileup, and then the average depth computed for 5kb windows across the genome.

*<u>MAT</u>* <u>locus determination</u>

To evaluate the mating type alleles present in each isolate, Illumina reads were aligned using bwa mem to a multifasta of both versions of the mating type locus (AF542529.2 and AF542528.2 (Lengeler et al., 2000b)). Depth at all positions was computed using Samtools mpileup, and then the average depth computed for the SXI1 and *STE20* genes for both idiomorphs. Nearly all isolates showed unique mapping to either the *MAT***a** or *MAT*α alleles of both genes; one isolate, Ftc158, showed significant mapping to both *MAT***a** and *MAT*α, though 2-fold more to *MAT*α. For the hybrid haploid isolates, the ancestry of the *MAT* locus was determined from the Structure site by site output.

#### Genome assembly and annotation

Illumina sequence for each isolate was assembled using Allpaths for 36 isolates (see Table S4 for release numbers for each assembly) or SPAdes 3.6.0 (with parameter–careful) for the remaining 3 isolates. Assemblies with both fragment and jump libraries were more contiguous than those with fragment only data (average of 84 or 561 scaffolds, respectively, **Table S4**). However there was little difference in the total contig length between assemblies with or without jump data (average 18.4Mb and 18.5Mb, respectively, **Table S4**).

The predicted protein coding gene set for each assembly was generated by combining three primary lines of evidence. Genes were transferred to each new assembly from the well annotated H99 assembly (Janbon et al., 2014) based on whole genome nucleotide alignments from nucmer. Genemark-ES (Ter-Hovhannisyan et al., 2008) was run on each assembly to generate a de novo set of calls. These two sets were combined and improved using PASA (Haas et al., 2008) with RNA-Seq data of three in vitro conditions (YPD, Limited media, and Pigeon guano) generated for H99 (Janbon et al., 2014) and for the VNB isolate Bt85 also input. Repetitive elements were removed from the gene set based on TransposonPSI (http://transposonpsi.sourceforge.net/) alignments or PFAM domains found only in transposable elements. The filtered set was assigned sequential locus identifiers across each scaffold. The average number of 6,944 predicted genes across all assemblies (**Table S4**) is close to the 6,962 predicted on the H99 reference.

#### Ortholog identification and comparison

To identify orthologs across the set of 45 *Cryptococcus* genomes (**Table S4**), proteins clustered based on BLASTP pairwise matches with expect< 1e-5 using ORTHOMCL v1.4 (Li et al., 2003). To identify orthologs specific to each of the serotype A lineages, we required that genes were present in 90% of the assembled genomes for VNI (36 or more) or VNB (8 or more) or all VNII (3 genomes). To confirm that orthologs were missing in the other two lineages, synteny was examined around each gene; in some cases this identified candidate orthologs missed by OrthoMCL, which were confirmed by BLASTP similarity and removed.

#### Phylogenetic analysis

A phylogeny for the sets of 159 or 164 isolates was inferred from SNP data using RAxML version 8.2.4 (Stamatakis, 2014) with model GTRCAT and 1,000 bootstrap replicates. A separate analysis of the phylogenetic relationship based on gene content included 40 *Cryptococcus neoformans* var. *grubii* serotype A genomes (28 VNI, 3 VNII, and 9 VNB), 1 *Cryptococcus neoformans* var. *neoformans* serotype D genome (JEC21), and 4 *C. gattii* genomes (WM276, R265, CA1873, and IND107) (**Table S4**). The total of 4616 single copy orthologs identified in all genomes were aligned individually with MUSCLE (Edgar, 2004) at the protein level, converted to the corresponding nucleotide sequence to maintain reading frame alignment, poorly aligning regions removed trimal (Capella-Gutiérrez et al., 2009), and invariant sites removed. A phylogeny was inferred using RAxML version 7.7.8 in rapid bootstrapping mode with model GTRCAT and 1,000 bootstrap replicates.

#### Population structure

To examine major population subdivisions, we examined how isolates clustered in a principal components analysis (PCA). SNP calls for all the isolates were compared using SMARTPCA (Patterson et al., 2006). To identify the major ancestry subdivisions and their contributions to the isolates appearing at intermediate positions in the PCA, a total of 338562 randomly subsampled positions containing variants in at least two isolates and less than 5% missing data were clustered using the Bayesian model-based program STRUCTURE v2.3 (Pritchard et al., 2000) in the site-by-site mode. Ancestry was plotted across the genome for each isolate using the maplotlib plotting package in Python.

For analysis of *Cryptococcus neoformans* var. *grubii* diploid isolates (**Table S3**), diagnostic SNPs for VNB and VNII were present exclusively in the respective group, and called for all VNB, VNII, and >=100 VNI isolates. Diagnostic SNPs for VNI were present exclusively in VNII and VNB, and called for all VNB, VNII, and >=100 VNI isolates.

Population genetic measures including Pi, Fst, and Tajima’s D were calculated using popGenome (Pfeifer et al., 2014). *d*_N_ and *d*_S_ measures were calculated from fixed SNPs in each lineage using codeml version 4.9c (Yang, 2007). To examine the distribution of the alleles within VNB, we first identified 445,193 alleles private to VNB (present in at least 1 VNB isolate and no VNI or VNII isolates). We subdivided VNB into four clades (VNBI-South America, VNBI-Africa, VNBII-South America, and VNBII-Africa) and calculated the number of those private alleles unique to each clade (present in that one clade and no others) and shared across VNB groups or geography (present in the two compared clades but no others). The Mantel test was conducted using the center-point of each country to determine distances between isolates and the number of SNPs between each pairwise set of isolates. The test was conducted using available Python software (https://github.com/jwcarr/MantelTest) with 1000 permutations and the upper tail test of positive correlation.

#### Linkage disequilibrium

Linkage disequilibrium was calculated in 500 bp windows of all chromosomes except for the ∼100kb mating type locus on chromosome 5 with vcftools version 1.14 (Danecek et al., 2011), using the --hap-r2 option with a minimum minor allele frequency of 0.1.

#### Population inference by fineStructure

Model-based clustering by fineStructure (Lawson et al., 2012) assigns individuals to populations based on a coancestry matrix created from SNP data, using either Markov chain Monte Carlo or stochastic optimisation. The algorithm uses chromosome painting, which is an efficient way of identifying important haplotype information from dense data, such as SNP data, and efficiently describes shared ancestry within a recombining population. Each individual is painted using all the other individuals as donors. For example, if an isolate *x* is clonal and a donor, the clonally related recipients will receive almost all of their genetic material from isolate *x*, and its closest relatives. This approach has been applied to analyze recombination in fungal (Engelthaler et al., 2014) and bacterial studies (Yahara et al., 2013).

fineStructure analysis (Lawson et al. 2012) was performed using an all lineage SNP matrix, with one representative of each clonal VNI population in order to infer recombination, population structure, and ancestral relationships of all lineages. A separate analysis of all VNI lineage isolates was also performed. This approach was based on the presence or absence of shared genomic haplotypes. ChromoPainter reduced the SNP matrix to a pairwise similarity matrix under the linked model, which utilises information on linkage disequilibrium, thus reducing the within-population variance of the coancestry matrix relative to the between-population variance. Since the MAT idiotypes introduce large bias into SNP analysis, they were removed to enable characterisation of more defined populations. There was no significant loss of sharing of genetic material when compared to retaining the MAT locus.

## Acknowledgments

We thank the Broad Institute Genomics Platform for generating DNA sequence for this study and Jose Munoz for helpful comments on the manuscript.

## Data access

All sequence data from this study have been submitted to GenBank under BioProject ID PRJNA 384983 (http://www.ncbi.nlm.nih.gov/bioproject); individual accession numbers are listed in Supplemental Tables S1 and S4.

## Author contributions

**Investigation**: JR, CAD, SMS, SS, CAC

**Validation**: JR, CAD, SMS, CAC

**Visualization**: JR, CAD, SMS, SS, CAC

**Writing – Original Draft Preparation**: JR, CAD, CAC

**Writing – Review & Editing**: JR, CAD, MCF, CAC, AA, MAB, DME, WM, FH, JMV, JH, AL, JP

**Resources**: MCF, CAC, MV, YC, JP, TB, TH, VP, ALC, AC, FH, MTI-Z, WM, DME, AA, JMV, JH

**Supervision**: CAC, MCF

**Funding Acquisition**: CAC, MCF

**Conceptualization**: CAC, MCF, AL

## Supplementary Figure Legends

Figure S1. Phylogenetic analysis of all sequenced isolates. Using a set of 876,121 SNPs across 165 isolates, the phylogenetic relationship was inferred with RAxML. The percentage of 1,000 bootstrap replicates that support each node is shown for major nodes with at least 90% support. Isolates with hybrid ancestry based on Structure are colored red.

Figure S2. Phylogenetic analysis of ancestry typed SNPs in hybrid isolates. SNPs with VNI and VNB ancestry were separated for each hybrid isolate, and combined with the wider set of SNPs for sequenced VNI or VNB isolates. Phylogenies were inferred using RAxML and the percentage of 1,000 bootstrap replicates that support each node is shown. The hybrid isolate in each phylogeny is highlighted with red text.

Figure S3. Loss of heterozygosity in AA diploid isolates. For each of the three diploid heterozygous isolates of AA ancestry, the frequency of heterozygous SNPs/kb within each isolate and the sequence depth is depicted for each of 14 chromosomes of H99. A. Bt66 has no apparent loss of heterozygosity. B. Cng9 shows a small LOH region at the start of scaffold 2, which also appears haploid in sequence depth. C. PMHc1045 shows more extensive LOH on multiple chromosomes, some of which are also associated with aneuploidy regions.

Figure S4. AD hybrids show high chromosomal aneuploidy. For each AD hybrid isolate the normalized depth of reads aligned to a combined AD reference is depicted across the chromosomes of the H99 and JEC21 genomes. In some isolates, the loss of a chromosome of one ancestry appears compensated by the gain of an extra copy in the other ancestry, noted by brackets, resulting in homozygosity of some regions of the hybrid genome.

Figure S5. Phylogenetic analysis of AD hybrid isolates. For each isolate, SNPs called against the H99 or JEC21 reference genome were separated, and those for H99 were combined with those from other VNB isolates. Phylogenies were inferred using RAxML.

Figure S6. Chromosomal aneuploidy highlighted by normalized read depth. The average depth of Illumina reads aligned to the 14 scaffolds of the H99 genome was plotted across the genome; whole and partial chromosomal aneuploidies were detected for 25 isolates.

Figure S7. Phylogenetic relationship and gene conservation for selected *Cryptococcus*. Using de novo assemblies and associated gene calls for the genome each isolate, orthologs were identified using OrthoMCL, and the aligned protein sequences of 4,616 single-copy genes were used to infer a phylogeny using RAxML. Bar graphs represent the numbers of core protein clusters shared across all genomes (green), conserved protein clusters found in a subset (grey), species specific protein clusters blue or orange) and clusters specific to a single genome (yellow).

Figure S8. Linkage disequilibrium as a function of distance expressed as the correlation coefficient r^2^ for a) the three lineages, VNB, VNII and VNI and b) South American and African VNB.

## Supplementary Tables Legends

Table S1. Metadata for sequenced isolates.

Table S2. Ancestry of Hybrid Strains from Structure

Table S3. Shared allele counts of diploid strains

Table S4. Genome assembly and annotation statistics.

Table S5. Lineage-specific genes.

Table S6. Genes conserved in C. *neoformans* var. *grubii* but absent in *C. gattii* (WM276, R265, CA1873, IND107) and C. *neoformans* var *neoformans* (JEC21)

Table S7. Ancestry for selected haploid isolates from Structure.

